# PathoBERT: A Hybrid Attention-Based Genomic Language Model for Read-Level Bacterial Pathogenicity Prediction

**DOI:** 10.64898/2026.09.27.754806

**Authors:** Salem A. El-aarag, Mario Flores, Mohamed E Hasan, Alaa E. Hemeida, Mahmoud ElHefnawi

## Abstract

**Motivation:** Although recent deep learning models have achieved promising results on read-level classification tasks, their robustness to realistic sequencing conditions, sensitivity to read length, and ability to support reliable genome-level inference remain incompletely characterized. Here, we present PathoBERT, a hybrid deep learning framework that integrates a LoRA-adapted DNABERT encoder with convolutional feature extraction, a modified Convolutional Block Attention Module (MCBAM), and Multi-Scale Convolutional Attention (MSCA) for bacterial pathogenicity prediction.

**Results:** The model demonstrated near-complete strand invariance and maintained robust performance under simulated sequencing errors, highlighting its suitability for real-world next-generation sequencing applications. The model operates on individual sequencing reads and supports genome-level inference through a read-aggregation strategy based on the Pathogenic Fraction (PathFrac), which combines read-level predictions using a majority-vote framework. At the read level, PathoBERT outperformed DeePaC at short and moderate fragment lengths (100–150 bp), achieving peak performance at 150 bp. At the genome level, PathoBERT achieved perfect separation between pathogenic and non-pathogenic genomes using PathFrac-based aggregation, resulting in perfect classification performance on the evaluated benchmark compared with competing approaches, namely DeePaC and PathogenFinder 2. Representation-level analyses further demonstrated that a progressive refinement of pathogen-associated features throughout the architecture, culminating in highly separable and biologically meaningful latent representations within the final attention-pooled embedding space. These findings demonstrate that integrating contextual genomic language models with attention-guided multi-scale feature extraction provides a robust framework for pathogenicity prediction from short-read sequencing data.

**Availability:** The supplementary data, datasets, and archived source code generated and analyzed during this study are available from the Zenodo repository (DOI: 10.5281/zenodo.11179933). The source code is available via GitHub at https://github.com/MahmoudElHefnawi/PathoBERT and https://github.com/salimalaarag/PathoBERT.

## Introduction

Bacterial pathogens evolve rapidly, and the vast diversity of bacteria, coupled with increasing human exposure, ensures the continued emergence of novel pathogens [1,2]. Distinguishing pathogenic from commensal bacteria remains challenging due to the complexity of the human microbiota [3]. Moreover, nonpathogenic bacteria can acquire pathogenic traits through horizontal gene transfer or mutations affecting virulence and antimicrobial resistance pathways, leading to the emergence of closely related pathogenic and nonpathogenic strains [4,5]. Consequently, rapid and reliable pathogen detection from next-generation sequencing data is essential, particularly given that most microbial diversity remains undiscovered and public sequence repositories continue to expand [6–8].

Pathogenicity prediction approaches can be broadly classified into protein content-based and read-based methods [9]. Protein content-based approaches rely on virulence factors or protein family profiles [10], enabling functional analyses and the discovery of novel virulence-associated proteins [11,12]. However, they depend on genome assembly and annotation and are less suitable for metagenomic applications. In contrast, read-based methods operate directly on raw sequencing reads [13], avoiding assembly requirements and facilitating faster metagenomic analyses [14,15]. These methods can be divided into taxonomic and non-taxonomic approaches. Taxonomic methods may fail to identify novel species when closely related references are unavailable [16], whereas non-taxonomic machine learning approaches can detect pathogenic potential without relying on taxonomic context [17]. Consequently, machine learning-based pathogenicity prediction has emerged as a promising alternative to reference-based methods for identifying novel pathogens [18]. More recently, deep learning has further advanced sequence-based classification capabilities [19].

Several machine learning and deep learning methods have been proposed for read-level pathogenicity prediction, including random forests, convolutional neural networks, recurrent neural networks, and reverse-complement architectures [20–22].

Deep learning models such as DeePaC have shown promising performance for read-level pathogenicity prediction using convolutional and recurrent architectures. However, these models are typically evaluated on fixed-length reads under controlled conditions, whereas real-world clinical and metagenomic sequencing data exhibit substantial variability in read length, orientation, and quality. Such heterogeneity poses challenges for model robustness and generalizability that remain insufficiently explored. Furthermore, read-level frameworks do not fully exploit genome-level inference strategies that aggregate noisy read-level predictions into stable organism-level decisions. This limitation is particularly relevant in metagenomic settings, where sequencing reads are fragmented and originate from complex microbial communities. In parallel, genome-level approaches such as PathogenFinder 2 [23] infer pathogenic potential from protein-and annotation-based genomic features, but their reliance on assembled genomes limits applicability to fragmented sequencing data.

To address these challenges, we developed a hybrid architecture that combines a pretrained DNABERT encoder with convolutional feature extraction, a modified Convolutional Block Attention Module (MCBAM), and Multi-Scale Convolutional Attention (MSCA) to capture both global contextual information and local sequence patterns. Low-Rank Adaptation (LoRA) was incorporated for efficient fine-tuning, while genome-level inference was enabled through a Pathogenic Fraction (PathFrac) aggregation framework based on majority voting of read-level predictions.

The proposed framework was evaluated across multiple dimensions relevant to real-world pathogen detection, including within-species and across-species generalization, varying read lengths, strand orientation, and sequencing-error robustness. Performance was benchmarked against DeePaC and PathogenFinder 2 at both read and genome levels. In addition, layer-wise representation analyses were conducted to characterize the discriminative properties of the learned embeddings.

Results showed that both PathoBERT and DeePaC were highly robust to strand orientation changes and sequencing noise. However, PathoBERT consistently outperformed DeePaC on shorter reads (100–150 bp), whereas DeePaC achieved superior performance on longer reads (250 bp). Genome-level analyses demonstrated that PathFrac-based aggregation enables accurate pathogenicity inference across whole genomes. Representation-level analyses further revealed that the MSCA module generated the most discriminative latent representations. Collectively, these findings highlight the value of combining contextual genomic language representations, multi-scale feature extraction, and hierarchical attention mechanisms for robust pathogen detection across diverse sequencing scenarios.

## Methods

### Dataset

Bacterial genomes were obtained from the IMG/M database on September 21, 2024 [24]. Human-associated pathogenic and non-pathogenic bacterial strains were identified using host and pathogenicity metadata and further validated through cross-referencing with external resources. Detailed genome-selection criteria, pathogenicity annotation procedures, exclusion rules, and database cross-validation are provided in Supplementary data 1, Section 1. To reduce class imbalance and minimize species-level sampling bias, a maximum of one strain per species was retained for the pathogenic class. The final dataset comprised 437 pathogenic (HP) and 77 non-pathogenic (non-HP) genomes. Genomic FASTA sequences were retrieved from NCBI using the corresponding BioProject accession numbers.

The dataset was partitioned into training, validation, and test subsets. The pathogenic class comprised 391, 8, and 14 genomes for training, validation, and testing, respectively, whereas the non-pathogenic class comprised 66, 4, and 5 genomes for the corresponding subsets.

### Preprocessing Stage

All nucleotide sequences were partitioned into fixed-length fragments of 150 bp. Sequences shorter than 150 bp were excluded, while longer sequences were segmented to ensure complete genome coverage. To mitigate class imbalance, non-pathogenic sequences were oversampled using multiple segmentation offsets, producing approximately 8 million fragments per class. Reverse-complement sequences were generated to promote strand-invariant learning.

To improve robustness to sequencing variability, paired-end synthetic reads were simulated from genomic FASTA sequences using the Mason simulator [25]. The final training dataset combined genomic fragments, reverse complements, and simulated reads for both classes and was organized into balanced stratified chunks. Detailed preprocessing procedures, oversampling strategy, fragment-generation workflow, and scalable read-simulation settings are provided in Supplementary data 1, Section 2 [26].

### Test Data Preparation

#### Intraspecies Test Set

To evaluate within-species generalization, unseen strains from species present in the training data were selected. For pathogenic species, held-out strains were reserved during dataset construction prior to training. For non-pathogenic species, additional strains from public repositories were retrieved and used exclusively for testing due to limited availability of internal held-out samples. All test strains were excluded from training and validation sets.

Model robustness was assessed using three variants of the intraspecies test set: (1) original sequences, (2) reverse-complement sequences, and (3) simulated Illumina-like reads generated using the Mason simulator. The dataset contained 455,671 non-overlapping 150 bp fragments for forward-strand evaluation, with reverse-complement counterparts used to test strand invariance. In addition, 500,000 paired-end reads were simulated under the same settings used during training.

To evaluate read-length effects, non-overlapping fragments of 100 bp, 150 bp, and 250 bp were generated, yielding 683,492, 455,671, and 273,405 fragments, respectively. Performance was compared with DeePaC across all length settings to assess robustness and generalization.

### Species-Level Test Set (Interspecies Evaluation)

Fourteen pathogenic species were initially designated for testing, but ten overlapped with DeePaC’s validation dataset. To avoid bias, these were removed, leaving four previously unseen pathogenic species. Overall, nine bacterial species (four pathogenic and five non-pathogenic) were used for out-of-distribution evaluation.

This benchmark included four novel pathogenic species and five non-pathogenic species, three of which (*Eubacterium ventriosum*, *Parabacteroides goldsteinii*, *Parabacteroides johnsonii*) were absent from both models’ training sets. Two *Bacteroides uniformis* strains overlapped with DeePaC’s validation data and are explicitly reported. Genome-level evaluation was performed consistently across all nine genomes, with prior exposure cases clearly indicated.

To assess read-length sensitivity, additional fragments of 100 bp, 150 bp, and 250 bp were generated, producing 420,243, 280,294, and 168,331 fragments, respectively. Benchmarking against DeePaC was conducted across all settings to evaluate robustness under varying read lengths.

### Model Architecture

A hybrid deep learning architecture (Figure 1) was developed by integrating a pretrained DNABERT [27–28] transformer encoder with convolutional and attention-based modules for bacterial pathogenicity classification. Input nucleotide sequences were tokenized and encoded using DNABERT. For parameter-efficient fine-tuning, Low-Rank Adaptation (LoRA) was applied to the query, key, and value projection matrices of the self-attention layers (rank = 16, scaling factor = 32, dropout = 0.05), while all pretrained DNABERT weights were frozen [29]. The resulting embeddings (B × L × H) were stripped of special tokens ([CLS], [SEP]) and transposed to B × H × L. A 1D convolutional layer (256 channels, kernel size 7), followed by batch normalization and GELU activation, was applied to capture local sequence motifs. Feature refinement was performed using a modified Convolutional Block Attention Module (MCBAM), which combines channel and spatial attention. Channel attention was computed using global average and max pooling, followed by a shared bottleneck network with two 1×1 convolutional layers and ReLU activation. Spatial attention was generated by concatenating channel-wise average and max-pooled features, followed by a 1D convolution and batch normalization [30–31].

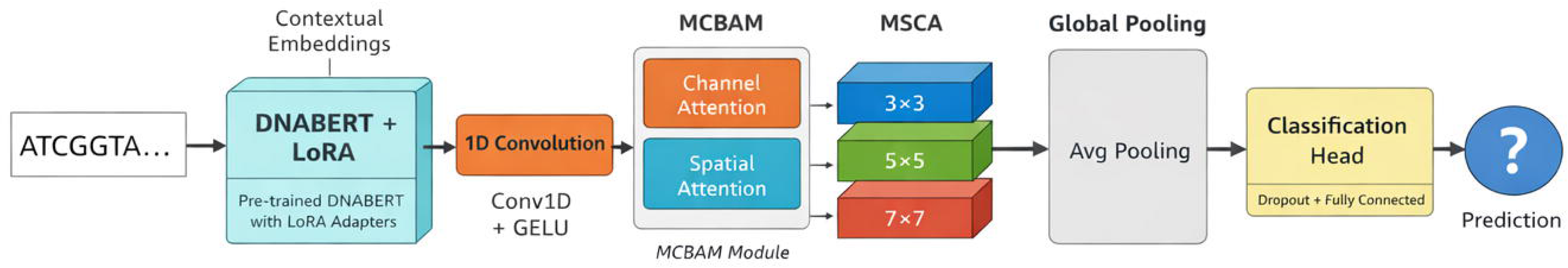

Multi-scale feature extraction was then performed using a Multi-Scale Convolutional Attention (MSCA) module, which applies parallel 1D convolutions with kernel sizes of 3, 5, and 7 [32]. The outputs are concatenated and refined using a channel attention mechanism based on global average pooling and a two-layer bottleneck network with sigmoid activation, followed by a 1×1 convolution for dimensional projection. The final representation was aggregated using adaptive average pooling and passed to a classification head consisting of dropout (0.3) and a fully connected linear layer for binary prediction. The model was optimized using AdamW (learning rate = 3 × 10⁻⁴, weight decay = 0.01) with BCEWithLogitsLoss.

Training was performed on a high-performance server equipped with 1024 GB system RAM and up to eight NVIDIA A100 GPUs, using a batch size of 256 per process. The distributed training framework allowed flexible multi-GPU execution depending on runtime resource allocation. Detailed Memory-Efficient Distributed Trainings are provided in Supplementary Data 1, Section 3.

### Evaluation metrics

#### Read-Level Pathogenicity Prediction

The proposed model was benchmarked against DeePaC, a widely used baseline for sequence-level pathogenicity prediction, using two independent datasets: an intraspecies test set (unseen strains from known species) and an interspecies test set (novel species). This setup enabled evaluation of both in-distribution performance and out-of-distribution generalization.

The model operates at the read level, assigning an independent pathogenicity probability to each sequencing read without requiring genome assembly, enabling real-time inference from high-throughput sequencing data.

Performance was evaluated using ten metrics. Discriminative ability was measured using AUROC and AUPR. Classification performance was assessed using accuracy, Matthews correlation coefficient (MCC), F1 score, precision, recall (sensitivity), false negative rate (FNR), and false positive rate (FPR). MCC was used as the primary balanced metric due to its robustness under class imbalance. Threshold-dependent metrics were computed using the decision threshold that maximized MCC on the validation set, while AUROC and AUPR were computed without thresholding. Results are reported separately for intraspecies and interspecies test sets.

### Robustness Testing Across Dataset Variants

Model robustness was evaluated on three intraspecies variants: original reads, reverse complements, and simulated Illumina-like reads generated using the Mason simulator. Performance uncertainty was estimated using a stratified bootstrap procedure (10,000 iterations), resampling positive and negative classes separately to preserve class balance and generate 95% confidence intervals for all metrics [33–35].

To compare conditions, a difference-based bootstrap was applied (5,000 iterations). For paired comparisons (original vs. reverse complement), a paired stratified bootstrap preserved read-level correspondence. For unpaired comparisons (original vs. simulated), independent stratified resampling was used. Metric differences were computed per iteration to obtain 95% confidence intervals for deltas.

For AUROC comparisons between paired conditions, the DeLong test for correlated ROC curves was applied, using the efficient implementation of Sun & Xu (2014), providing variance estimates, Z-scores, and p-values [36–37]. Effect sizes for proportion-based metrics were quantified using Cohen’s h, while Wasserstein distance was used to assess distributional shifts between original, reverse complement, and simulated reads [38–40].

### Impact of Fragment Length

Model performance was additionally evaluated across multiple fragment length settings for both intraspecies and interspecies test sets to assess robustness to read length variation.

### Genome-Level Pathogenicity Prediction via Read Aggregation

Both the proposed model and DeePaC operate at the read level rather than directly performing genome-level inference. Each genome is first fragmented into sequencing reads, which are independently processed by the model to generate per-read pathogenicity probabilities. These read-level predictions are subsequently aggregated to produce a genome-level decision.

Let p = [p_1_, p_2_,…, p_n_] denote the set of pathogenicity probabilities assigned to n reads derived from a single genome. Genome-level inference is computed using two complementary summary statistics.

The mean pathogenicity score is defined as the average probability across all reads:

Mean score = (sum of pᵢ for i = 1 to n) / n

The pathogenic fraction (PathFrac) is defined as the proportion of reads classified as pathogenic:

PathFrac = (number of reads with pᵢ > 0.5) / n

Genome-level classification is then determined using a majority-vote rule:

GenomePred = 1 if PathFrac > 0.5, otherwise GenomePred = 0

GenomePred = 1 if Mean pathogenicity score > 0.5, otherwise 0

Thus, a genome is classified as pathogenic when the majority of its reads are predicted to be pathogenic. This aggregation scheme converts noisy read-level predictions into a stable genome-level decision by integrating evidence across multiple independent observations, thereby improving robustness to stochastic variability in individual read predictions.

In contrast, PathogenFinder 2 is a genome-level classifier that directly estimates pathogenic potential from assembled genome sequences using protein- and annotation-derived features, without intermediate read-level inference. The model was evaluated using default parameters across all assembled genome sequences in the test dataset. PathogenFinder 2 was executed using the default settings of the publicly available web server (https://genepi.food.dtu.dk/pathogenfinder) on all assembled genome sequences included in the test dataset.

### Layer-Wise Contribution to Class Separation Analysis

To interpret how the model distinguishes pathogenic from non-pathogenic sequences, a layer-wise representation analysis was conducted under both interspecies and intraspecies settings [41–42]. This dual evaluation assesses whether hierarchical feature learning remains consistent across different levels of genomic similarity and distributional shift.

Feature embeddings were extracted from four representational stages: DNABERT + LoRA outputs, CNN features, MCBAM-refined representations, and the final MSCA with global pooling. These stages capture the progressive transformation from contextual sequence embeddings to task-specific latent representations. Analysis was conducted from three complementary perspectives: statistical and geometric separability, distributional and clustering structure, and task-aligned linear separability.

### Statistical and Geometric Separability

Class separation was quantified using the Fisher Score and Normalized Fisher Score, measuring the ratio of inter-class to intra-class variance, where higher values indicate stronger discriminative structure [43–46]. Geometric separation was further assessed using cosine separation, defined as the average cosine distance between class centroids, capturing angular discrimination independent of magnitude [47]. These measures were computed for each layer across both evaluation settings to track how separability evolves through the network.

### Distributional and Clustering Structure

Distributional divergence between pathogenic and non-pathogenic embeddings was measured using Wasserstein distance at each layer, quantifying shifts between class distributions [48]. Clustering quality was assessed using silhouette scores, evaluating intra-class compactness and inter-class separation to characterize latent space organization across the network hierarchy [49–50].

### Task-Aligned Linear Separability

To assess how explicitly pathogenicity information is encoded, linear probe analysis was performed at each layer. A logistic regression classifier was trained on frozen embeddings, and probe accuracy was used as a measure of linear separability. This provides a direct estimate of task-relevant information content across representational stages [51–52].

## Results and Discussion

### 1. Read-Level Pathogenicity Prediction

#### 1.1 Robustness Testing of PathoBERT Across Intraspecies test set variants

We evaluated ten performance metrics on three Intraspecies test set variants: (1) original sequences, (2) reverse complements, and (3) simulated reads with Illumina-like error profiles using mason simulator. As shown in Figure 2, point estimate results of performance metrics for the reverse complement sequences as (AUROC: 0.9420, MCC: 0.7308) were nearly identical to those obtained from the original sequences (AUROC: 0.9417, MCC: 0.7274). However, the simulated reads results as (AUROC: 0.9390, MCC: 0.7214) exhibited only a limited reduction in performance relative to the original sequences. All three conditions achieve AUROC > 0.93, indicating strong discriminative ability between positive and negative classes. The near-equivalence of AUROC and AUPR values (difference < 0.002) suggests balanced class distribution or symmetric performance.

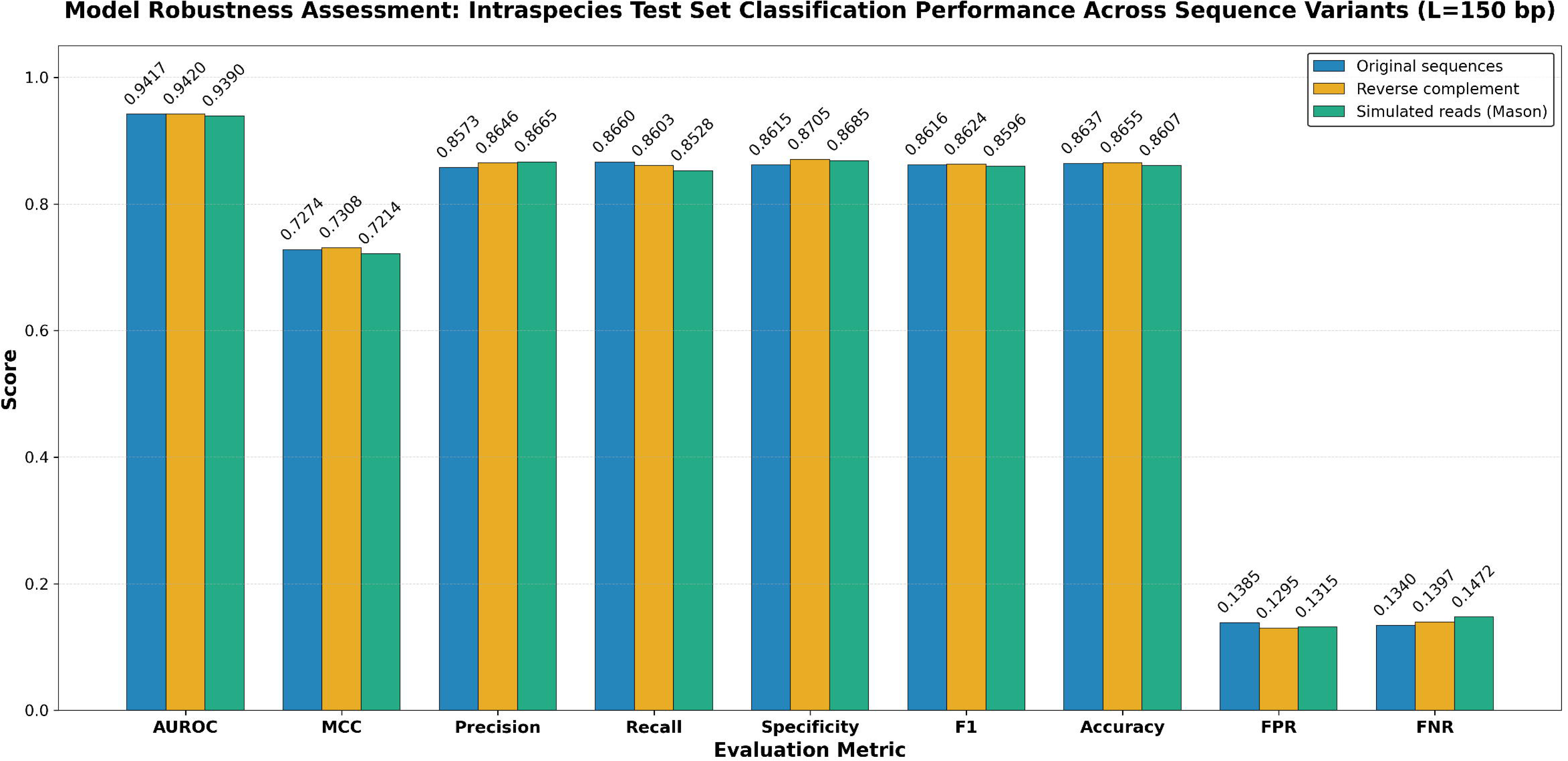

Narrow Bootstraping confidence intervals estimates for ten performance metrics (max width = 0.004) across the three conditions indicates high statistical stability and reliability.

**The reverse-complement** results showed overlapping confidence intervals for AUROC, AUPR, MCC, F1, and accuracy with the original sequences results, although the optimal classification threshold increased slightly from 0.5226 to 0.5377 (+0.0151). This indicates no significant differences, confirming strong strand invariance. The DeLong test confirmed that there was no statistically significant difference in AUROC between the original and reverse-complement sequences. Although the reverse-complement evaluation produced a slightly higher AUROC (0.9420 vs. 0.9417), the observed improvement was extremely small (ΔAUROC = +0.0003) and statistically non-significant (Z = 1.39, p = 0.164). 95% confidence intervals for metric differences (Reverse Complement − Original Sequences) showed no statistically significant differences were observed for AUROC, AUPR, and F1 since the confidence intervals for these metrics all included zero. Although several point estimates showed slight positive shifts, the bootstrap analysis suggests that these variations are likely attributable to sampling variability rather than a systematic performance effect. While the confidence intervals for the remaining metrics did not cross zero, indicating statistically detectable differences, though the absolute magnitudes of these changes were extremely small (|Δ| ≤ 0.009 with the largest CI bound reaching 0.01). The Wasserstein distance between the reverse complement and original sequence predictions was extremely small (Wasserstein distance: 0.00036; normalized: 0.00095), indicating virtually no distributional shift. Effect size analysis using Cohen’s h revealed minimal practical differences between (Reverse Complement − Original Sequences) across all metrics (range: 0.0002–0.0264), confirming that the observed performance changes, while statistically detectable in some cases, lack practical import.

**The simulated reads** results showed overlapping confidence intervals only for AUPR and F1 with the original sequences results, although the optimal classification threshold increased slightly from 0.5226 to 0.5477 (+0.0251). 95% confidence intervals for metric differences (Simulated Sequences − Original Sequences) showed no statistically significant differences were observed for AUPR since the confidence intervals for this metric included zero. While the confidence intervals for the remaining metrics did not cross zero, indicating statistically detectable differences, though the absolute magnitudes of these changes were extremely small (|Δ| ≤ 0.0131 with the largest CI bound reaching 0.0152). Simulated sequences showed small shift in prediction distributions compared with the original sequences (Wasserstein distance: 0.00797; normalized: 0.02092), suggesting that sequencing-error simulation slightly alters the prediction distribution while preserving overall model performance. Effect size analysis using Cohen’s h revealed minimal practical differences between ( Simulated Sequences − Original Sequences) across all metrics (range: 0.0006–0.037), confirming that the observed performance changes, while statistically detectable in some cases, lack practical import.

Taken together, the near-identical performance across original and reverse-complement sequences confirms that the model learns biologically meaningful strand-independent representations, while the limited degradation observed for Mason-simulated reads suggests practical applicability to real-world sequencing and metagenomic environments. Detailed bootstrap confidence intervals, pairwise confidence-interval comparisons, DeLong tests, Wasserstein distances, and effect-size analyses are provided in **Supplementary Data 2.xlsx (PathoBERT_Robustness, Tables S1–S9)**. Visualizations of metric differences with corresponding 95% confidence intervals are available in the **Forest_plot_PathoBERT** worksheet.

#### 1.2 Robustness Testing of DeePaC Across Intraspecies test set variants

As a benchmark comparison, we assessed **DeePaC**, a leading deep learning model for pathogenic sequence classification, using the same **Intraspecies test sets (L = 150 bp)**. As shown in Figure 3, performance on reverse-complement sequences was virtually indistinguishable from that obtained on the original sequences, yielding **AUROC** and **MCC** values of **0.8979** and **0.6117**, respectively, compared with **0.8979** and **0.6109** for the original data. Evaluation on simulated reads resulted in a slight performance decrease (**AUROC = 0.8939, MCC = 0.6014**), indicating only a minor sensitivity to sequencing-related variations. Across all three conditions, **AUROC values remained above 0.89**, although lower than that achieved by PathoBERT. In addition, the close agreement between AUROC and AUPR values, with differences below **0.0037**, suggests a balanced class distribution and uniform predictive performance across classes.

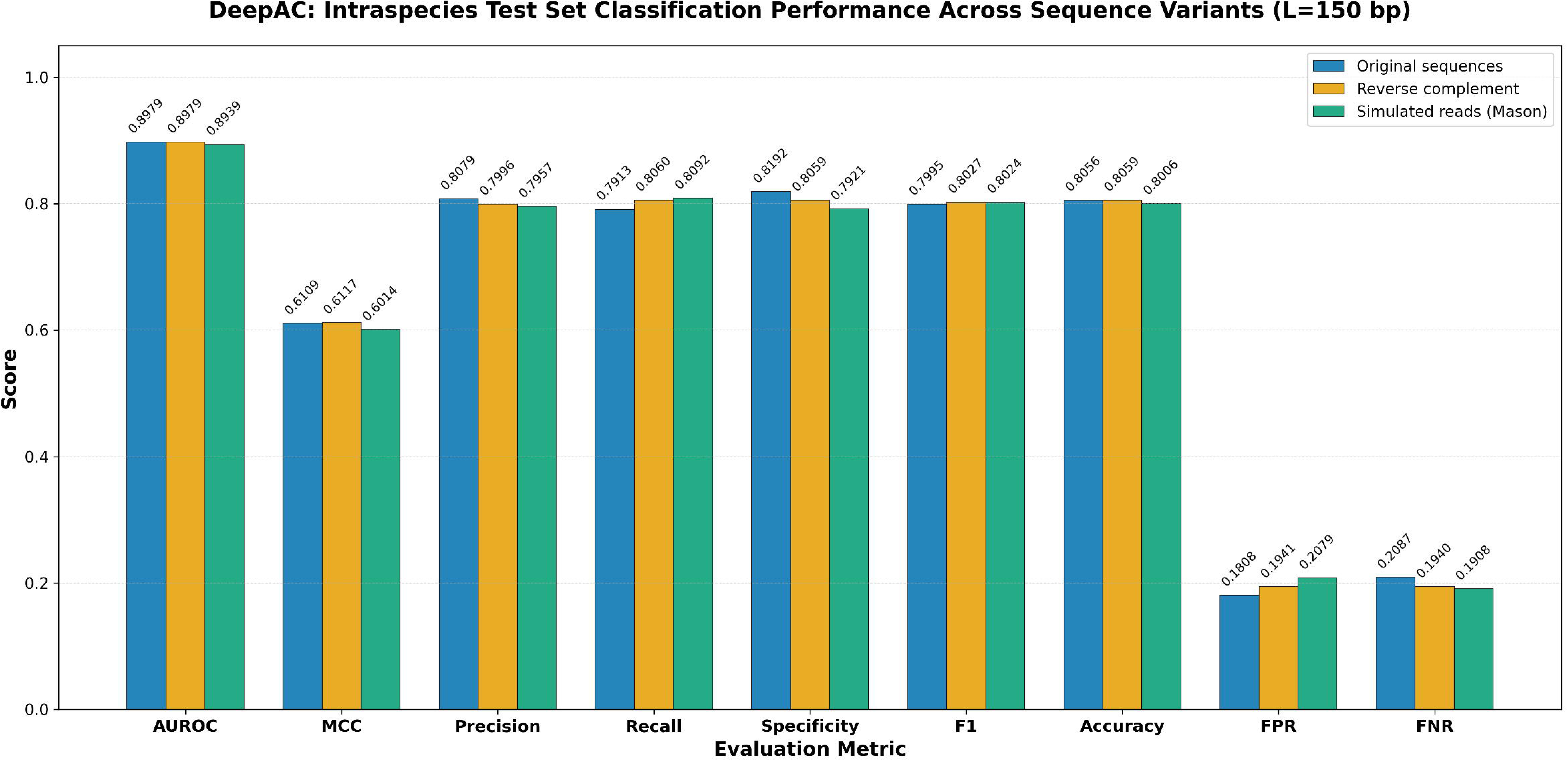

Bootstrap-derived confidence intervals were consistently narrow across all evaluated metrics, with a maximum width of 0.0046, indicating high precision and reliability of the performance estimates. These findings suggest that DeePaC exhibits stable predictive performance across original sequences, reverse complements, and simulated reads.

**The reverse-complement** evaluation yielded confidence intervals for AUROC, AUPR, MCC, F1 score, and accuracy that overlapped with those obtained for the original sequences, mirroring the pattern observed for PathoBERT. Although the optimal classification threshold decreased slightly from 0.6131 to 0.5980 (Δ = −0.0151), overall predictive performance remained essentially unchanged, supporting strong strand invariance. Consistent with this observation, the DeLong test detected no statistically significant difference in AUROC between the original and reverse-complement datasets (Z = 0.259, p = 0.795). Further analysis of the 95% confidence intervals for performance differences (Reverse Complement − Original Sequences) showed that the intervals for AUROC, AUPR, and F1 score included zero, indicating no statistically significant differences. While the confidence intervals for the remaining metrics excluded zero, suggesting statistically detectable differences, the magnitude of these effects was negligible (|Δ| < 0.0145, with the largest confidence interval bound reaching only 0.0165). Similarly, the prediction distributions were nearly indistinguishable, as evidenced by an extremely small Wasserstein distance (0.0006; normalized = 0.00186), indicating virtually no distributional shift between the two conditions. Effect size analysis using Cohen’s h further demonstrated trivial practical differences across all evaluated metrics (range: 0.0001–0.0334). Collectively, these findings indicate that reverse complementation has a negligible impact on DeePaC performance, with any observed differences being statistically minor and lacking practical significance. **For the simulated reads,** only the confidence intervals for AUPR overlapped with those obtained from the original sequences, although the optimal classification threshold decreased slightly from 0.6131 to 0.5879 (Δ = −0.0252). Analysis of the 95% confidence intervals for performance differences (Simulated Reads − Original Sequences) indicated no statistically significant differences for AUPR, F1 score, and accuracy, as the corresponding confidence intervals included zero. Although the confidence intervals for the remaining metrics excluded zero, suggesting statistically detectable differences, the magnitude of these changes was minimal (|Δ| ≤ 0.0206, with the largest confidence interval bound reaching only 0.0226). The simulated reads exhibited a small shift in prediction distributions relative to the original sequences, as reflected by a low Wasserstein distance (0.0057; normalized = 0.0178). This finding suggests that sequencing-error simulation introduces only minor alterations to the prediction distribution while largely preserving overall classification performance. Consistent with this observation, effect size analysis using Cohen’s h revealed trivial practical differences across all evaluated metrics (range: 0.0006–0.0370), indicating that the observed performance changes, although statistically detectable in some cases, have negligible practical significance. Overall, DeePaC demonstrated robust and consistent performance across all sequence variants. The reverse-complement results indicate that the model does not rely on strand-specific sequence patterns and exhibits strong reverse-complement invariance. Likewise, the simulated-read analysis demonstrates that DeePaC retains its predictive capability in the presence of sequencing-like perturbations introduced by Mason simulation. Detailed results, including bootstrap confidence intervals, pairwise confidence-interval comparisons, DeLong tests, Wasserstein distance analyses, and effect-size estimates, are provided in **Supplementary data 2.xlsx (DeePaC_Robustness, Tables S11–S19).** Corresponding visualizations of metric differences with 95% confidence intervals are presented in the **Forest_plot_DeePaC worksheet.**

#### 1.3 Impact of Fragment Length on Intraspecies Test Set Classification Performance

We evaluated eleven performance metrics across three sequence fragment lengths (L = 100, 150, and 250 bp) on an intraspecies test set (Table 1, Figure 4). At L = 100, our model substantially outperformed DeePaC across all metrics, achieving an AUROC of 0.8555 (vs. 0.8032), MCC of 0.5444 (vs. 0.4354), and F1 score of 0.7662 (vs. 0.7185), with notably lower false positive rates (0.2171 vs. 0.3008). At L = 150, our model demonstrated optimal and highly robust performance, attaining an AUROC of 0.9417, AUPR of 0.9409, MCC of 0.7274, and accuracy of 0.8637, while DeePaC achieved AUROC of 0.8979, AUPR of 0.8985, and MCC of 0.6109 at the same length. This represents a substantial performance gain for our model from L = 100 to L = 150 (ΔAUROC = +0.0862, ΔMCC = +0.1830), indicating that moderate fragment lengths provide sufficient discriminative signal. Interestingly, DeePaC shows steady improvement across all lengths, with AUROC increasing from 0.8032 (L=100) to 0.8979 (L=150) to 0.9661 (L=250). Notably, at L=250, DeePaC surpasses our proposed model achieving an AUROC of 0.9661, MCC of 0.7879, and AUPR of 0.9666, compared to 0.9428, 0.7297, and 0.9420 for our model, respectively, indicating that DeePaC benefits more from longer sequence context. This crossover suggests fundamental architectural differences between the two models: our model appears to capture relevant sequence features efficiently at shorter to moderate lengths (L ≤ 150) and exhibits diminishing returns beyond this point, whereas DeePaC — consistent with its deep convolutional design — benefits more substantially from extended sequence contexts. These results suggest that our model’s multi-scale attention mechanism is particularly effective at extracting discriminative features from shorter sequences, while DeePaC requires longer fragments to achieve optimal performance.

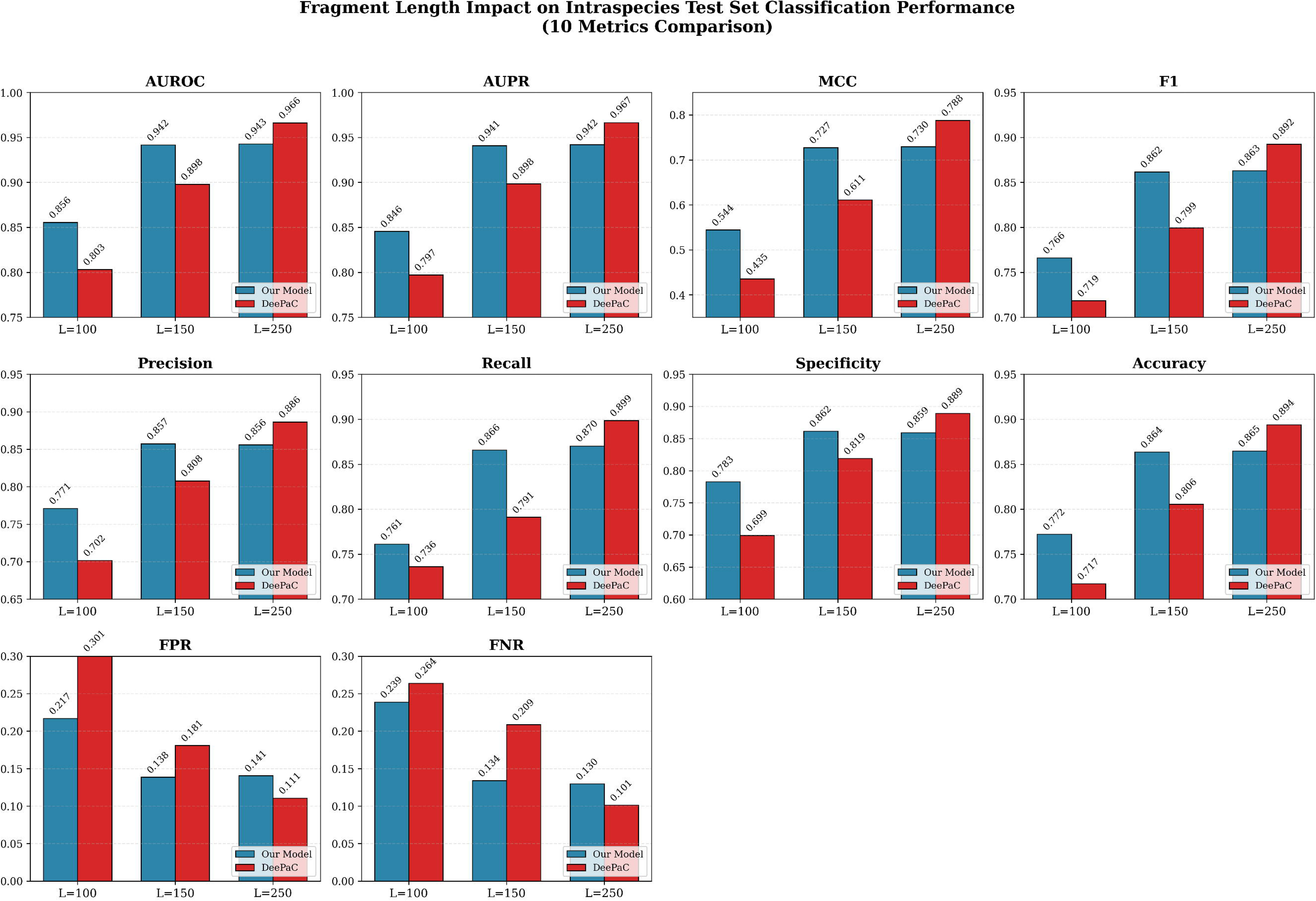

**Table 1:** Comparison of classification performance between the proposed PathoBERT model and DeePaC on the intraspecies test set across three sequence fragment lengths (L = 100, 150, and 250 bp).. Best results for each fragment length are shown in bold.

| Metric | L = 100 |  | L = 150 |  | L = 250 |  |
| --- | --- | --- | --- | --- | --- | --- |
|  | PathoBERT | DeePaC | PathoBERT | DeePaC | PathoBERT | DeePaC |
| AUROC | **0.8555** | 0.8032 | **0.9417** | 0.8979 | 0.9428 | **0.9661** |
| AUPR | **0.8457** | 0.7971 | **0.9409** | 0.8985 | 0.9420 | **0.9666** |
| MCC | **0.5444** | 0.4354 | **0.7274** | 0.6109 | 0.7297 | **0.7879** |
| F1 score | **0.7662** | 0.7185 | **0.8616** | 0.7995 | 0.8632 | **0.8925** |
| Precision | **0.7712** | 0.7016 | **0.8573** | 0.8079 | 0.8561 | **0.8864** |
| Recall | **0.7612** | 0.7361 | **0.8660** | 0.7913 | 0.8704 | **0.8987** |
| Specificity | **0.7829** | 0.6992 | **0.8615** | 0.8192 | 0.8594 | **0.8893** |
| Accuracy | **0.7723** | 0.7173 | **0.8637** | 0.8056 | 0.8648 | **0.8939** |
| FPR | **0.2171** | 0.3008 | **0.1385** | 0.1808 | 0.1406 | **0.1107** |
| FNR | **0.2388** | 0.2639 | **0.1340** | 0.2087 | 0.1296 | **0.1013** |

#### 1.5 Impact of Fragment Length on Species-Level Classification

For read-based species-level classification, the test set necessarily included all available species due to the fixed data partitioning used during model evaluation. Among these, two nonpathogenic Bacteroides uniformis strains were present in DeePaC’s original validation set, conferring prior exposure for DeePaC on these strains. This biases the specificity comparison in DeePaC’s favor for these two species. Results are reported with this caveat; the primary conclusions regarding read-length robustness and generalization remain unaffected as these analyses focus on different aspects of model behavior.

We evaluated our proposed model on species-level taxonomic classification across three fragment lengths (L=100, 150, and 250 bp) and compared its performance against DeePaC (Table 2, Figure 5). At L = 100, our model substantially outperformed DeePaC across all metrics, achieving an AUROC of 0.8740 (vs. 0.8102), MCC of 0.5927 (vs. 0.4737), and F1 score of 0.7999 (vs. 0.7675), with notably lower false positive rates (0.1652 vs. 0.3246). At L = 150, our model demonstrated optimal and highly robust performance, attaining an AUROC of 0.9480, AUPR of 0.9630, and MCC of 0.7677, while DeePaC achieved AUROC of 0.9209, AUPR of 0.9329, and MCC of 0.6786 at the same length. This represents a substantial performance gain for our model from L = 100 to L = 150 (ΔAUROC = +0.0740, ΔMCC = +0.1750), indicating that moderate fragment lengths provide sufficient discriminative signal. DeePaC shows steady improvement across all lengths, with AUROC increasing from 0.8102 (L=100) to 0.9209 (L=150) to 0.9814 (L=250). Notably, at L=250, DeePaC surpasses our model, achieving an AUROC of 0.9814, MCC of 0.8547, and AUPR of 0.9854, compared to 0.9478, 0.7685, and 0.9631 for our model, respectively, indicating that DeePaC benefits more from longer sequence context. This crossover suggests fundamental architectural differences: our model captures relevant sequence features efficiently at shorter to moderate lengths (L ≤ 150) with diminishing returns beyond this point, whereas DeePaC — consistent with its deep convolutional design —benefits substantially from extended sequence contexts. Our model’s multi-scale attention mechanism appears particularly effective at extracting discriminative features from shorter sequences, while DeePaC requires longer fragments to achieve optimal performance.

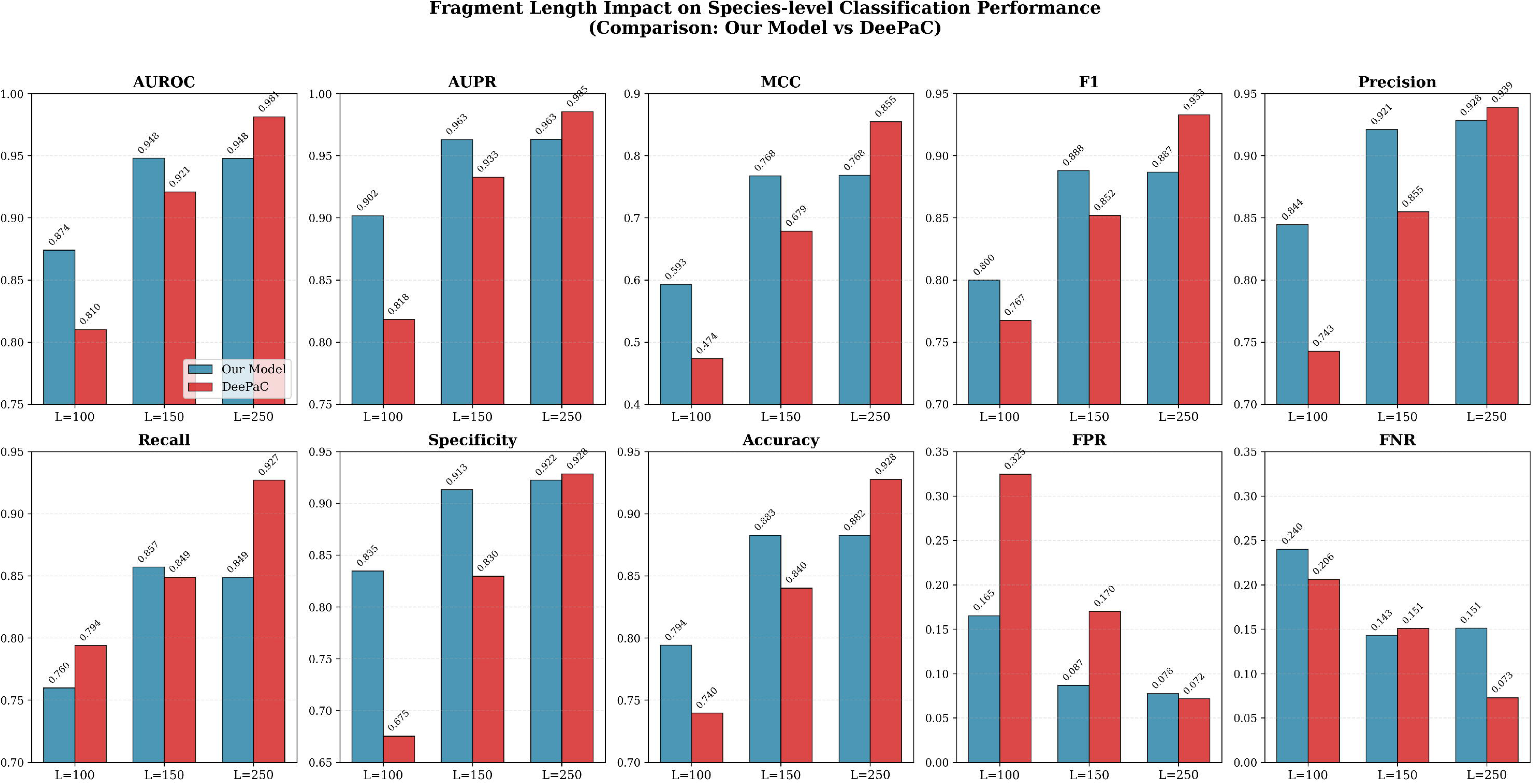

**Table 2:** Comparison of classification performance between the proposed PathoBERT model and DeePaC on the Species-level test set across three sequence fragment lengths (L = 100, 150, and 250 bp). Best results for each fragment length are shown in bold.

| Metric | L = 100 bp |  | L = 150 bp |  | L = 250 bp |  |
| --- | --- | --- | --- | --- | --- | --- |
|  | PathoBERT | DeePaC | PathoBERT | DeePaC | PathoBERT | DeePaC |
| AUROC | **0.8740** | 0.8102 | **0.9480** | 0.9209 | 0.9478 | **0.9814** |
| AUPR | **0.9018** | 0.8183 | **0.9630** | 0.9329 | 0.9631 | **0.9854** |
| MCC | **0.5927** | 0.4737 | **0.7677** | 0.6786 | 0.7685 | **0.8547** |
| F1 score | **0.7999** | 0.7675 | **0.8879** | 0.8520 | 0.8867 | **0.9329** |
| Precision | **0.8444** | 0.7427 | **0.9211** | 0.8549 | 0.9283 | **0.9388** |
| Recall | 0.7598 | **0.7940** | **0.8571** | 0.8491 | 0.8488 | **0.9271** |
| Specificity | **0.8348** | 0.6754 | **0.9133** | 0.8298 | 0.9224 | **0.9285** |
| Accuracy | **0.7942** | 0.7396 | **0.8828** | 0.8403 | 0.8825 | **0.9277** |
| FPR | **0.1652** | 0.3246 | **0.0867** | 0.1702 | 0.0776 | **0.0715** |
| FNR | 0.2402 | **0.2060** | **0.1429** | 0.1509 | 0.1512 | **0.0729** |

From a practical bioinformatics perspective, our model offers a superior trade-off between sensitivity and specificity at L = 150 (recall = 0.8571, specificity = 0.9133), making it suitable for applications where both false positives and false negatives are costly, such as clinical strain typing or outbreak surveillance. Conversely, DeePaC at L = 250 may be preferred when maximum accuracy is required and longer reads are available. These findings underscore that fragment length is not merely a technical parameter but a critical determinant of model performance, and optimal length should be empirically determined based on both model architecture and specific application requirements.

### 2. Genome-Level Pathogenicity Prediction via Read Aggregation

We compared our proposed model against PathogenFinder 2 and two DeePaC variants (150 bp and 250 bp) using both mean prediction scores and PathFrac—read-level predictions were aggregated using the majority-vote framework described in Section 4.2.

The proposed model demonstrated clear separation between pathogenic and nonpathogenic genomes, assigning low mean prediction scores to all nonpathogenic genomes (0.2686–0.3908) and substantially higher scores to all pathogenic genomes (0.8392–0.8964). In contrast, both DeePaC configurations correctly identified all pathogenic genomes but misclassified *Eubacterium ventriosum* as pathogenic with mean scores 0.5865 and 0.5873 for 150 bp and 250 bp respectively. Notably, increasing fragment length from 150 bp to 250 bp improved pathogenic confidence scores for DeePaC on true pathogenic genomes but did not resolve this false-positive prediction. PathogenFinder 2 exhibited a distinct prediction profile. While it correctly classified several genomes with high confidence, it produced false-positive predictions (0.8781) for *Bacteroides uniformis* strains and generated an ampigous score (0.5176) for *Streptomyces somaliensis* (Table 3, Supplementary data 3, Figures S3- S8).

**Table 3.** Genome-level pathogenicity prediction scores for benchmark genomes.

| Genome | PathogenFinder 2* |  | Proposed Model |  | DeePaC (150 bp) |  | DeePaC (250 bp) |  |
| --- | --- | --- | --- | --- | --- | --- | --- | --- |
|  | Mean | prediction | Mean | PathFrac | Mean | PathFrac | Mean | PathFrac |
| **Nonpathogenic genomes** |  |  |  |  |  |  |  |  |
| *Eubacterium ventriosum* ATCC 27560 |  | 0.1719 | 0.3908 | 0.3464 | **0.5865** | **0.6397** | **0.5873** | **0.6385** |
| *Bacteroides uniformis* 82G1 |  | **0.8781** | 0.2890 | 0.1722 | 0.3268 | 0.2085 | 0.2685 | 0.1518 |
| *Parabacteroides goldsteinii* DSM 19448 |  | 0.3777 | 0.2686 | 0.1538 | 0.3159 | 0.2058 | 0.2462 | 0.1424 |
| *Parabacteroides johnsonii* DSM 18315 |  | 0.2263 | 0.2719 | 0.1594 | 0.3170 | 0.2006 | 0.2491 | 0.1398 |
| **Pathogenic genomes** |  |  |  |  |  |  |  |  |
| *Nocardia veterana* DSM 44445 |  | 0.9434 | 0.8392 | 0.8910 | 0.8503 | 0.9438 | 0.9629 | 0.9907 |
| *Streptomyces somaliensis* DSM 40738 |  | 0.5176 | 0.8964 | 0.9336 | 0.8389 | 0.9273 | 0.9676 | 0.9893 |
| *Klebsiella variicola* WUSM KV_02 |  | 0.9767 | 0.8921 | 0.8910 | 0.8101 | 0.8980 | 0.9360 | 0.9758 |
| *Klebsiella quasipneumoniae* WUSM |  | 0.9797 | 0.8848 | 0.9267 | 0.7993 | 0.8878 | 0.9283 | 0.9702 |

\* Pathogen Finder 2 reports only mean prediction (no PathFrac available).

#### Comparison between mean prediction and PathFrac

For our model, PathFrac values generally aligned with mean predictions but offered clearer separation. For nonpathogenic strains, PathFrac tended to be slightly lower than the mean prediction, for example, *E. ventriosum* showed a mean prediction of 0.3908 but a PathFrac of 0.3464 (clearly nonpathogenic under a 0.5 threshold). Similarly, for pathogenic strains, PathFrac tended to be slightly higher than the mean prediction, reinforcing classification confidence. Similarly, for DeePaC models, PathFrac exceeded mean predictions in nearly all cases, for pathogenic genomes, suggesting that aggregating sequence-level scores into PathFrac amplifies true pathogenic signals while suppressing noise—though this same effect produced false positives for nonpathogenic *E. ventriosum*. For nonpathogenic strains, PathFrac tended to be slightly lower than the mean prediction except for E. ventriosum as it was assigned as pathogenics by DeePaC models. DeePaC exhibited progressively higher pathogenic fractions with increased fragment length, reaching near-complete pathogenic consensus at 250 bp (0.9893–0.9907). However, this increased confidence does not translate into improved discrimination for challenging negative cases, where PathoBERT demonstrates superior specificity. Collectively, these PathFrac distributions demonstrate that PathoBERT achieves superior genome-level classification not through inflated pathogenic confidence, but through improved calibration around the majority-vote threshold. Specifically, our model maintains clear separation between nonpathogenic genomes (PathFrac < 0.35) and pathogenic genomes (PathFrac > 0.89), creating a wide decision gap of more than 0.54 between classes. This separation substantially reduces susceptibility to threshold-sensitive errors and highlights the effectiveness of the read aggregation framework for robust genome-level pathogenicity inference.

The read-to-genome aggregation strategy employed by both the proposed model and DeePaC offers several advantages: (1) it naturally handles variable coverage depths, (2) it can detect mixed or low-abundance populations, and (3) it provides statistical confidence through read-level replicates. The read-aggregation approach inherently produces more stable genome-level decisions than single-value genome-level classifiers, as the ensemble of thousands of read predictions reduces variance and provides a natural buffer against ambiguous cases. It captures intra-genomic heterogeneity while enforcing a majority-vote constraint, thereby reducing sensitivity to sporadic misclassification at the read level and improving interpretability of genome-scale pathogenicity inference.

These findings underscore that the PathFrac metric — the proportion of reads exceeding the 0.5 pathogenicity threshold — proved to be a stable and discriminative aggregate for genome-level classification in PathoBERT.

The relatively small genome-level benchmark reflects the limited availability of bacterial genomes that satisfy stringent inclusion criteria, including unambiguous pathogenicity annotation, species-level label consistency, and independence from prior benchmark exposure. Given that pathogenicity is evaluated at the species level, these results should be regarded as initial evidence supporting improved species-level pathogenicity discrimination rather than definitive proof of broad generalization.

#### Layer-Wise Contribution To Class Separation

Understanding why certain deep learning architectures succeed or fail in bacterial pathogenicity prediction requires examining their internal representations across multiple analytical lenses. A layer-wise representation analysis was performed across both the interspecies and intraspecies separately (Table 4). MSCA + Pooling consistently outperforms all other architectures across nearly all evaluated metrics, exhibiting a distinctly superior and stable separability profile. Its Fisher Scores (interspecies: 560.08; intraspecies: 498.89) are one to three orders of magnitude higher than those of CNN, MCBAM, and BERT + LoRA, indicating an exceptionally well-disentangled feature space with extreme inter-class dispersion relative to intra-class variance. This trend is consistently supported by normalized Fisher and cosine separation metrics, where MSCA + Pooling maintains near-maximal angular discrimination while preserving compact class structure, in contrast to CNN and MCBAM which achieve moderate but substantially lower separability, and BERT + LoRA which shows comparatively weak class alignment in the embedding space. The superiority of MSCA + Pooling is further reinforced by distributional structure analysis. The Silhouette Score (0.4181 for both interspecies and intraspecies datasets)—substantially higher than all competing layers—confirms that the learned representations form compact, well-separated clusters with strong intra-class cohesion and clear inter-class boundaries. CNN and MCBAM exhibit moderate clustering structure, with significantly lower Silhouette values, while BERT + LoRA shows near-degenerate clustering behavior, reflecting weak intrinsic class separability. This hierarchy is further corroborated by Wasserstein distance, where MSCA + Pooling achieves the highest inter-class distributional separation (0.0301 interspecies; 0.0290 intraspecies), compared with markedly smaller values for CNN, MCBAM, and BERT + LoRA, indicating progressively increasing overlap in their learned feature distributions.

**Table 4.** Layer-wise representation separability across interspecies and intraspecies evaluation datasets.

| **Metric** | **Layer** | **Interspecies** | **Intraspecies** |
| --- | --- | --- | --- |
| **Fisher Score** | BERT + LoRA | 0.1175 | 0.0394 |
|  | CNN | 87.0798 | 15.9816 |
|  | MCBAM | 51.8654 | 13.8521 |
|  | **MSCA + Pooling** | **560.0804** | **498.8863** |
| **Normalized Fisher Score** | BERT + LoRA | 0.1761 | 0.0467 |
|  | CNN | 79.2662 | 16.2484 |
|  | MCBAM | 49.3675 | 13.2560 |
|  | **MSCA + Pooling** | **536.9915** | **505.3735** |
| **Cosine Separation** | BERT + LoRA | 0.5419 | 0.5100 |
|  | CNN | 0.9561 | 0.8303 |
|  | **MCBAM** | **0.9600** | 0.8612 |
|  | MSCA + Pooling | 0.9559 | **0.9005** |
| **Wasserstein Distance** | BERT + LoRA | 0.0003 | 0.0002 |
|  | CNN | 0.0125 | 0.0039 |
|  | MCBAM | 0.0049 | 0.0024 |
|  | **MSCA + Pooling** | **0.0301** | **0.0290** |
| **Silhouette Score** | BERT + LoRA | 0.0100 | 0.0013 |
|  | CNN | 0.1255 | 0.0232 |
|  | MCBAM | 0.0849 | 0.0849 |
|  | **MSCA + Pooling** | **0.4181** | **0.4181** |
| **Linear Probe Accuracy** | BERT + LoRA | 0.5102 | 0.5100 |
|  | CNN | 0.8941 | 0.8303 |
|  | **MCBAM** | **0.9079** | 0.8612 |
|  | MSCA + Pooling | 0.9024 | **0.9005** |

Notably, the near-identical performance of MSCA + Pooling across interspecies and intraspecies evaluations suggests that the model learns a generalizable and taxonomically robust representation, maintaining strong separability even under reduced phylogenetic distance.

Importantly, task-level evaluation reveals a critical divergence between geometric separability and downstream linear discriminability. Although MSCA + Pooling achieves the highest structural separability overall (Table 4, Figure 6), its Linear Probe Accuracy (0.9024 interspecies; 0.9005 intraspecies) is only marginally higher than MCBAM and comparable to CNN. This discrepancy suggests a saturation effect, where extreme embedding separability does not proportionally translate into improved linear decision boundaries. In contrast, MCBAM achieves competitive linear probe performance with substantially lower Fisher Scores, indicating more efficient alignment between representation geometry and linear classification objectives.

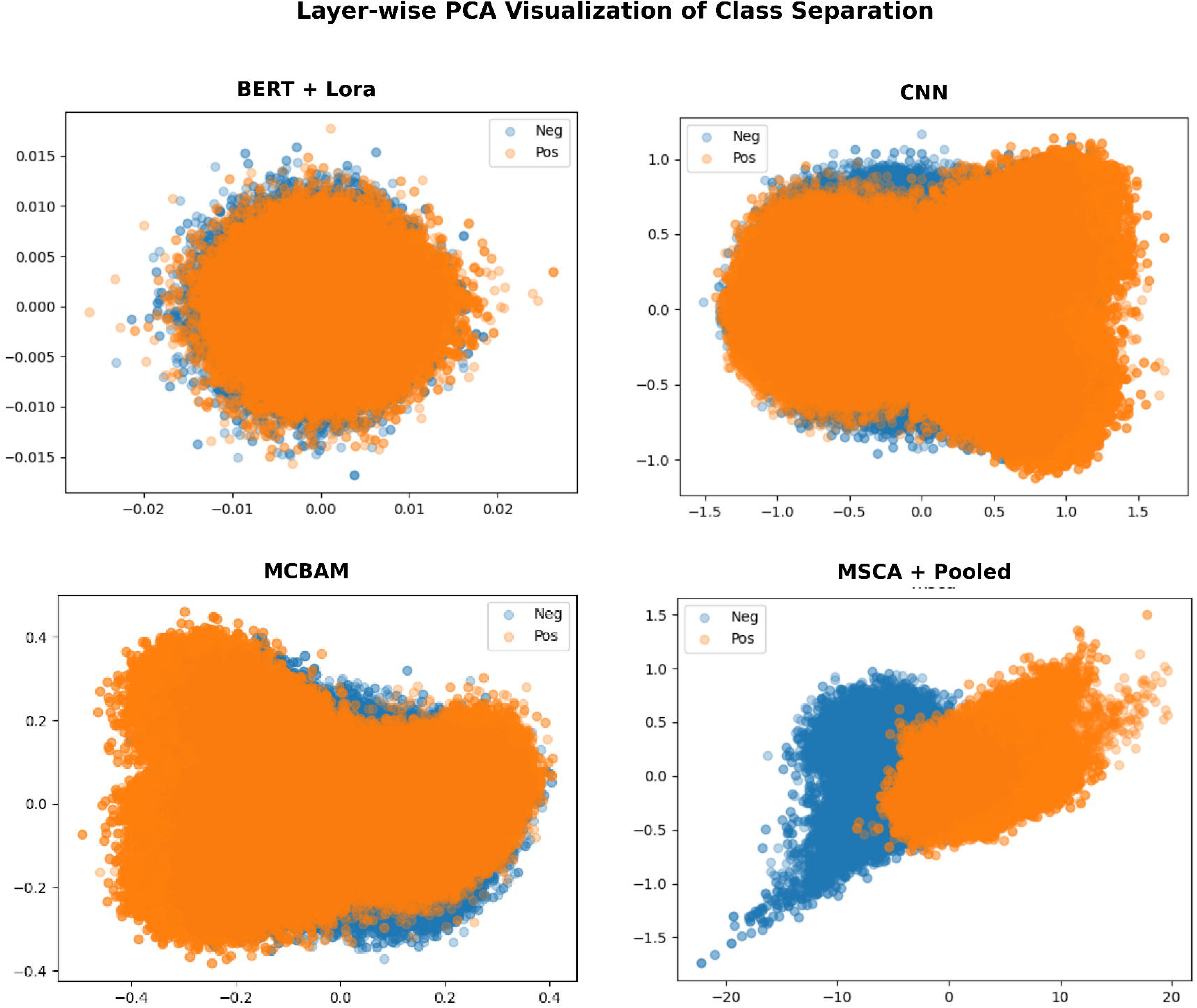

## Conclusion

This study demonstrates that our hybrid architecture model PathoBERT provides robust, accurate, and interpretable pathogenicity prediction across both read- and genome-level analyses. At the read level, the model exhibits robustness to sequence orientation and sequencing-error perturbations while consistently outperforming DeePaC at practical short-read lengths, highlighting its ability to capture informative local and contextual genomic patterns under realistic sequencing conditions. At the genome level, aggregation of read-level predictions through the PathFrac framework substantially improves classification stability between pathogenic and nonpathogenic genomes within the evaluated benchmark for PathoBERT. While PathoBERT maintains superior calibration and specificity relative to competing approaches namely DeepPac and PathogenFinder 2.

Beyond predictive performance, Representation-level analysis further reveals that the proposed MSCA + Pooling module induces highly structured and strongly separable latent spaces, as evidenced by Fisher score, cosine separation, Wasserstein distance, and silhouette analysis. However, linear probe evaluation exposes a key insight: extreme geometric separability does not necessarily translate into proportional gains in linear discriminability, indicating a saturation effect in representation utility.

Collectively, these findings demonstrate that integrating contextual embeddings with hierarchical convolutional and attention-based refinement enables the transformation of raw genomic sequences into robust pathogenicity-aware representations. The proposed framework therefore offers a promising foundation for short-read pathogen detection and genomic risk assessment.

Taken together, these findings demonstrate that accurate pathogenicity prediction benefits from a combination of biologically informed sequence modeling, multi-scale feature extraction, and robust aggregation strategies. Our framework provides a scalable and generalizable solution for genome-wide pathogen detection, with strong potential for real-world applications in clinical diagnostics, metagenomic surveillance, and microbial risk assessment.

Future work will focus on improving calibration under extreme distribution shifts, reducing redundancy in over-separated embedding spaces, extending the framework to multi-class microbial phenotype prediction (opportunistic, pathogens, nonpathogens), reducing redundancy in over-separated embedding spaces, adaptation to long-read sequencing platforms with variable read lengths, integration with attention-based interpretability methods to identify virulence-related sequence motifs, and prospective validation on real-time clinical metagenomic samples to assess operational performance in diagnostic settings.

## Acknowledgements

The authors gratefully acknowledge Prof. Mario Flores and his laboratory, Department of Biomedical Engineering, University of Texas at San Antonio, for generously providing computational server resources that were instrumental in conducting this study. The authors are also grateful to the National Research Centre computational cloud of Prof Mahmoud ElHefnawi and to the super computer of the library of Alexandria.

## Author contributions

S.A.E and M.E conceived and designed the study. S.A.E and M.E developed the methodology. S.A.E performed Data acquisition, Model development, Traininig, evaluation, performed formal analysis, and prepared the initial manuscript. S.A.E and M.F run the scripts. M.E and M.F revised the paper. All authors agreed to the published version of the manuscript.

## Data availability

The The supplementary data cited in this paper, datasets and source code generated and analyzed during the current study are available in Zenodo repository: https://doi.org/10.5281/zenodo.11179933. <u>The source code is also available at</u> https://github.com/MahmoudElHefnawi/PathoBERT <u>or</u> https://github.com/salimalaarag/PathoBERT.

---

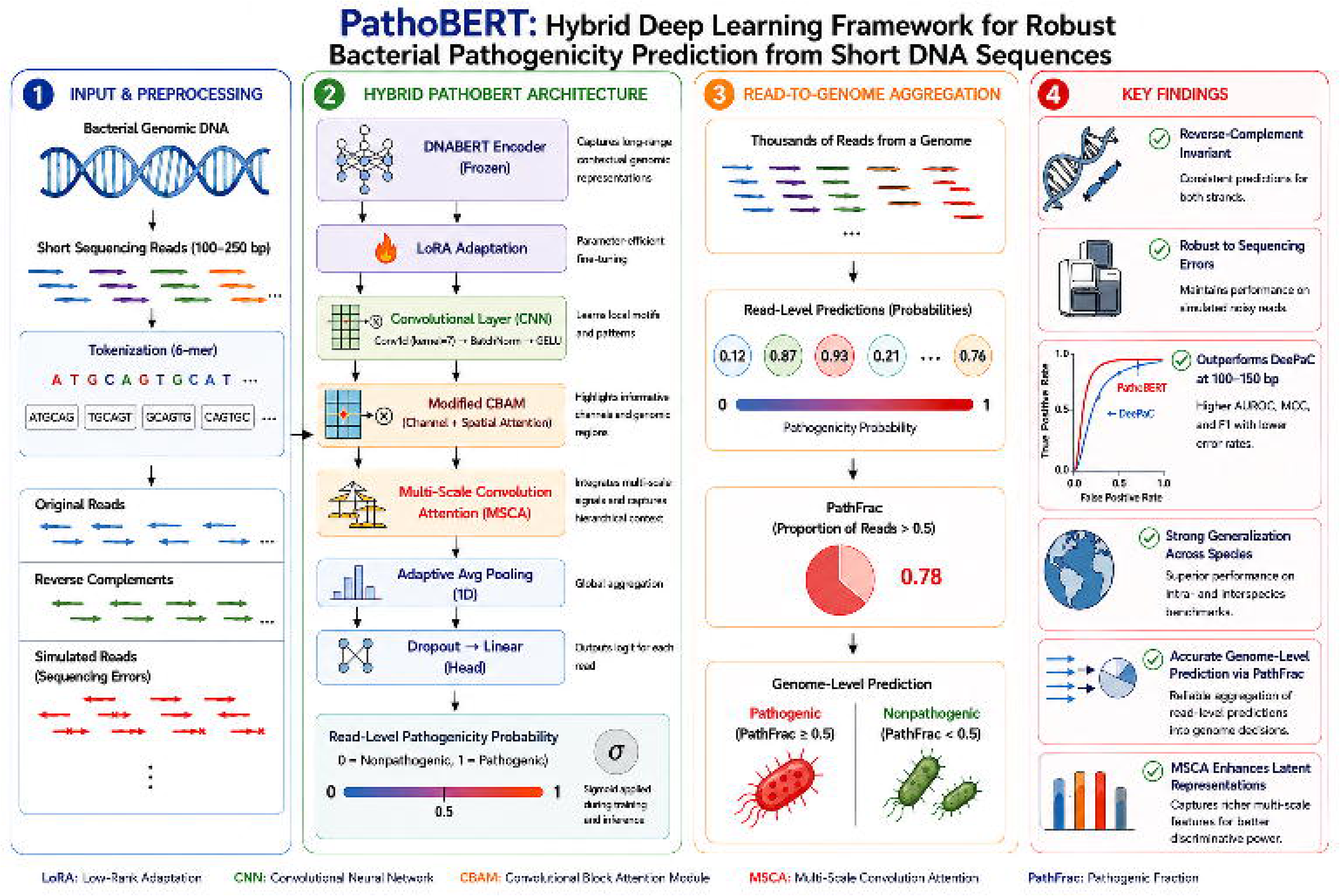

