## Supplementary data 1 for "PathoBERT: A Hybrid Attention-Based Genomic Language Model for Read-Level Bacterial Pathogenicity Prediction": Supplementary data 1.docx

**1. Dataset**

On September 21, 2024, we accessed the IMG/M database and downloaded a dataset containing all relevant bacterial genome information. As a preliminary step, we filtered the data to include only entries classified as Bacteria. To identify bacteria with human hosts, we applied specific criteria: records containing "human" or "Homo sapiens" in the "Host Name," "Ecosystem Category," or "Habitat" fields were selected. For pathogenic classification, we searched for records with the keyword "Pathogen" in the fields "Phenotype," "Relevance," or "Specific Ecosystem," or where there were disease-related entries in the "Diseases" field. Non-pathogens were identified by searching for the term "Non-pathogen" in the "Phenotype" field. Additionally, we cross-referenced certain strains with other databases such as KEGG or PATRIC to validate their pathogenic classification. We excluded nonpathogenic strains of species known to include pathogenic variants, such as Escherichia coli, Campylobacter jejuni, Listeria monocytogenes, Staphylococcus epidermidis, Staphylococcus aureus, Francisella tularensis, Clostridium difficile, andClostridium botulinum, from the non-pathogenic list. In cases where reference sequences of pathogenic strains had been removed from the NCBI database due to quality issues, we sought to identify these strains in PATRIC or find alternative strains from the same species using the same pathogenic classification criteria. Given that the number of pathogenic strains exceeded the number of non-pathogenic strains, we faced an issue of imbalanced data. This can adversely affect machine learning algorithms, potentially leading to suboptimal classification results when there is a large discrepancy in sample size ratios. To mitigate this imbalance, we restricted our dataset to one strain per species, resulting in a final list of 437 (409) pathogenic (HP) and 77 non-pathogenic (non-HP) strains. The genomic FASTA sequence files were retrieved from NCBI by querying with the BioProject accession numbers provided in the IMG dataset. The dataset was partitioned into training, validation, and test subsets. The pathogenic class comprised 391 genomes for training, 8 genomes for validation, and 14 genomes for testing. The non-pathogenic class consisted of 66 genomes for training, 4 genomes for validation, and 5 genomes for testing.

### **2. Preprocessing Stage**

All nucleotide sequences were partitioned into fixed-length, non-overlapping fragments of 150 bp. Terminal sequence regions shorter than 150 bp were adjusted by realigning them to the final 150 bases of the parent sequence, thereby preserving complete sequence coverage while avoiding the creation of undersized fragments. Sequences with lengths below 150 bp were discarded. For longer sequences, contiguous fragments were generated using a deterministic sliding-window approach with a stride equal to the fragment length.

To address class imbalance, non-pathogenic sequences were oversampled to achieve approximate parity with the number of pathogenic fragments (approximately 8 million reads per class). This oversampling strategy involved segmenting each sequence using multiple starting offsets (0, 25, 50, 75, 100, and 125 bp), which produced overlapping fragments without introducing synthetically generated sequence content. In addition, reverse-complement counterparts were generated for all fragments to facilitate orientation-invariant feature learning.

**Scalable read simulation**

We developed bash script that implements a sophisticated pipeline for simulating Illumina sequencing reads from a reference genome. It's designed to handle large genomes by splitting them into manageable chunks, processing them in parallel, and generating paired-end reads with realistic characteristics.

Prior to simulation, FASTA headers were standardized and genome sequences were partitioned into segments of up to 200 kb to facilitate efficient large-scale processing and parallel execution. This segmentation strategy reduced memory requirements and enabled balanced distribution of simulation workloads across multiple compute cores using GNU Parallel [1]. Contigs shorter than the minimum fragment-length requirement were excluded. Read simulation was performed independently for each sequence segment using Mason [2] with a read length of 150 bp, a mean fragment size of 350 bp, and a fragment-size standard deviation of 50 bp to augment the training data and mimic realistic Illumina sequencing characteristics. invalid or insufficiently long sequence segments were automatically excluded from processing. The filtering threshold (500bp) is derived from the fragment size distribution, ensuring realistic simulation while optimizing compute resources. The total number of simulated reads was allocated proportionally to sequence length, ensuring that larger genomes contributed a correspondingly greater number of reads while preserving the overall genomic composition of the dataset.

Following completion of all simulation jobs, generated read files were merged into consolidated paired-end datasets for downstream analysis.

This workflow enabled efficient simulation of millions of reads from large genome collections while maintaining realistic sequencing characteristics and proportional genome representation.

**The final training dataset** combined original fragments, reverse complements and synthetic paired-end reads for each class. Data were organised into balanced stratified chunks.

### **3. Memory-Efficient Distributed Training**

To enable scalable training on large genomic datasets, all input samples were stored as pre-tokenized NumPy arrays and accessed using memory-mapped loading, thereby avoiding full in-memory dataset materialization. A custom iterable dataset was implemented to support PyTorch DistributedDataParallel, with deterministic partitioning of samples across distributed processes. Dataset lengths were truncated to ensure equal sample allocation across ranks, while additional sharding was applied at the data-loader worker level to prevent overlap between worker processes.

Training data were organized into disk-resident chunks that were loaded sequentially during training. This chunk-based strategy enabled efficient processing of datasets exceeding GPU memory capacity while maintaining deterministic sample ordering and reproducibility through epoch-dependent shuffling.

Memory optimization strategies were incorporated at multiple stages of training. After each optimization step, gradients were cleared using optimizer.zero_grad(set_to_none=True), an intermediate tensors (including batch data, logits, loss values, and prediction probabilities) after each training iteration were explicitly deleted, and release of chunk-level resources through removal of data loaders and dataset objects following chunk completion. GPU memory cache was subsequently cleared using torch.cuda.empty_cache(), while host memory was reclaimed through periodic invocation of garbage collection (gc.collect()).

To improve data throughput, pinned host memory and data prefetching were employed within the data-loading pipeline, reducing transfer overhead between CPU and GPU memory. Worker-level deterministic initialization ensured reproducible parallel loading across epochs.

**Refrences**

1. Tange, O. (2011). GNU Parallel - The Command-Line Power Tool. USENIX ;login: The USENIX Magazine, 36(1), 42–47.
2. Holtgrewe M. (2010) Mason—a read simulator for second generation sequencing data. Technical Report FU Berlin.
