## supplementary data 3 for "PathoBERT: A Hybrid Attention-Based Genomic Language Model for Read-Level Bacterial Pathogenicity Prediction": Supplementary data 3.docx

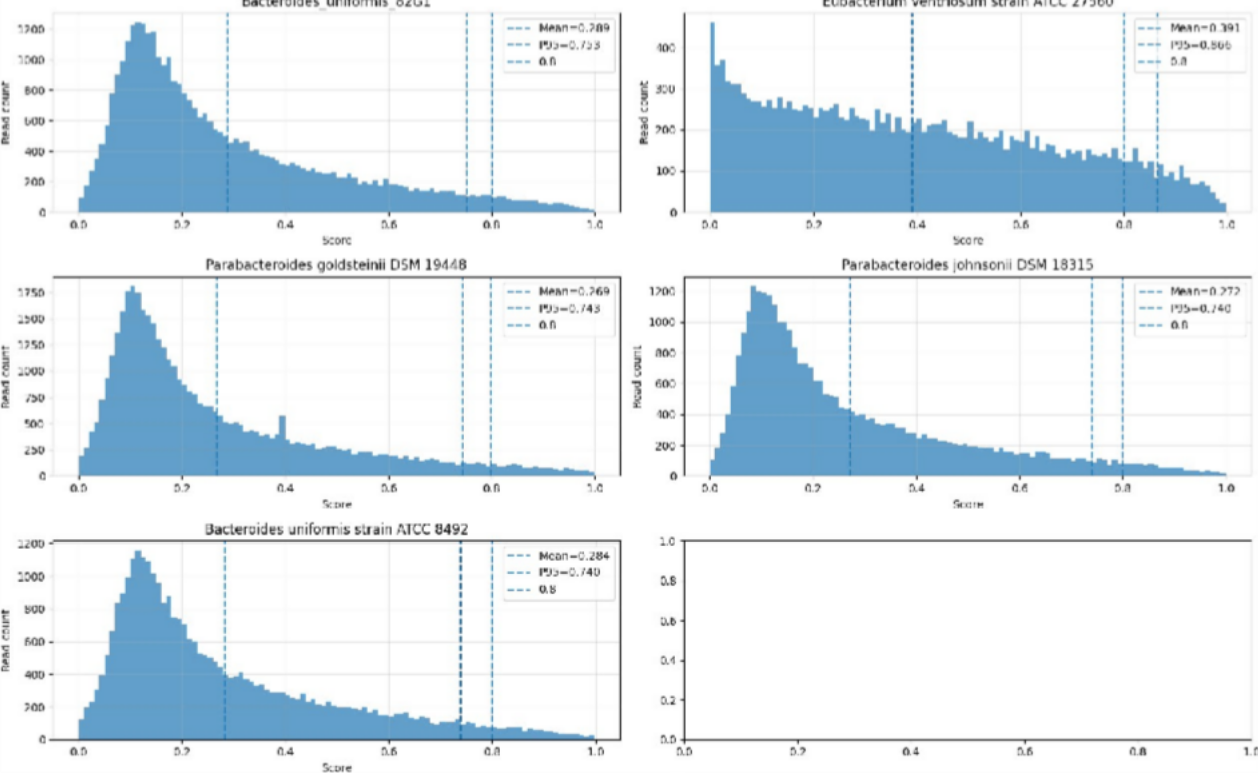

**Figure S3. Read-level pathogenicity score distributions for nonpathogenic genomes.** Per-read scores from the proposed model are shown for five commensal strains. Distributions are left-skewed, with most reads scoring below the 0.5 pathogenic threshold (vertical dashed line). Embedded bar charts indicate the pathogenic fraction (PathFrac, red) and nonpathogenic fraction (NonPathFrac, gray) of reads. All PathFrac values range from 0.15 to 0.35, below the 0.5 genome-level decision boundary, resulting in correct nonpathogenic classifications for all five genomes.

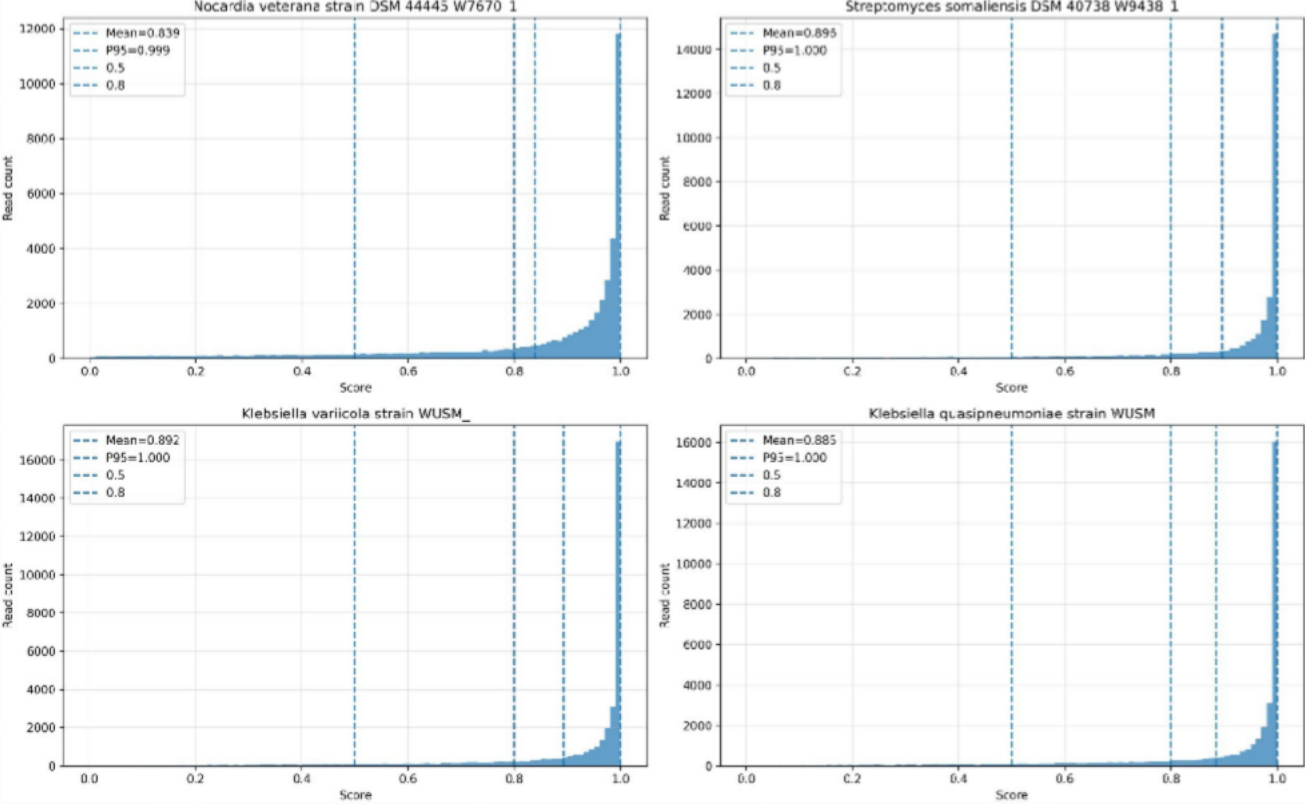

**Figure S4. Read-level pathogenicity score distributions for pathogenic genomes predicted by the proposed model.** Per-read scores from the proposed model are shown for four pathogenic strains: *Nocardia veterana* DSM 44445, *Streptomyces somaliensis* DSM 40738, *Klebsiella variicola* WUSM, and *Klebsiella quasipneumoniae* WUSM. Distributions are right-skewed, with most reads scoring above the 0.5 pathogenic threshold (vertical dashed line). Embedded bar charts indicate the pathogenic fraction (PathFrac, red) and nonpathogenic fraction (NonPathFrac, gray) of reads. All PathFrac values range from 0.89 to 0.93, above the

0.5 genome-level decision boundary, resulting in correct pathogenic classifications for all four genomes.

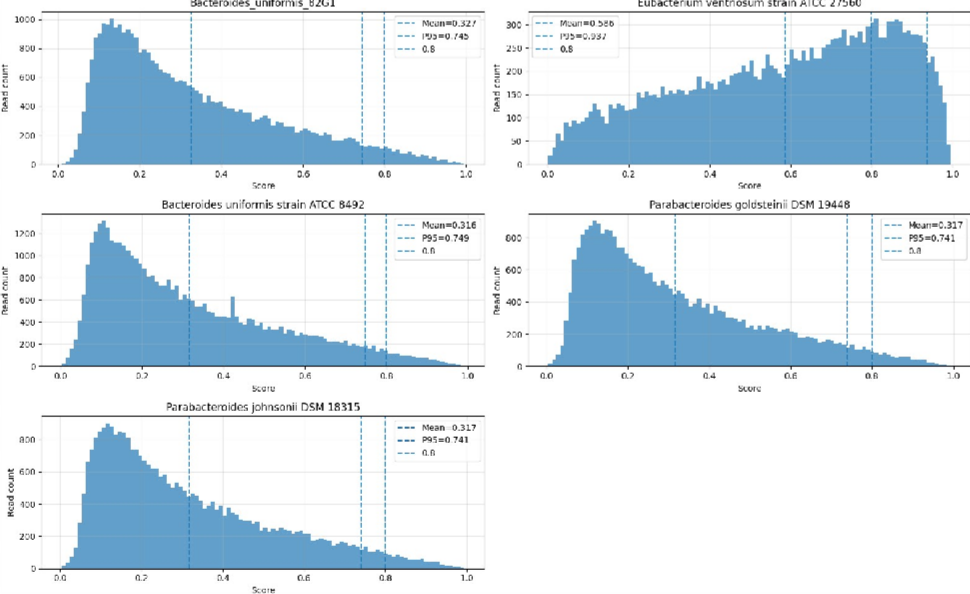

**Figure S5. DeePaC (150 bp) read-level predictions for nonpathogenic genomes.** Four of five commensal strains show left-skewed score distributions with PathFrac values below 0.5 (dashed line), yielding correct nonpathogenic calls. *Eubacterium ventriosum* ATCC 27560 is an outlier: its PathFrac of 0.6397 exceeds the

0.5 threshold, driving a false-positive pathogenic prediction. Embedded bar charts show PathFrac (red) versus NonPathFrac (gray).

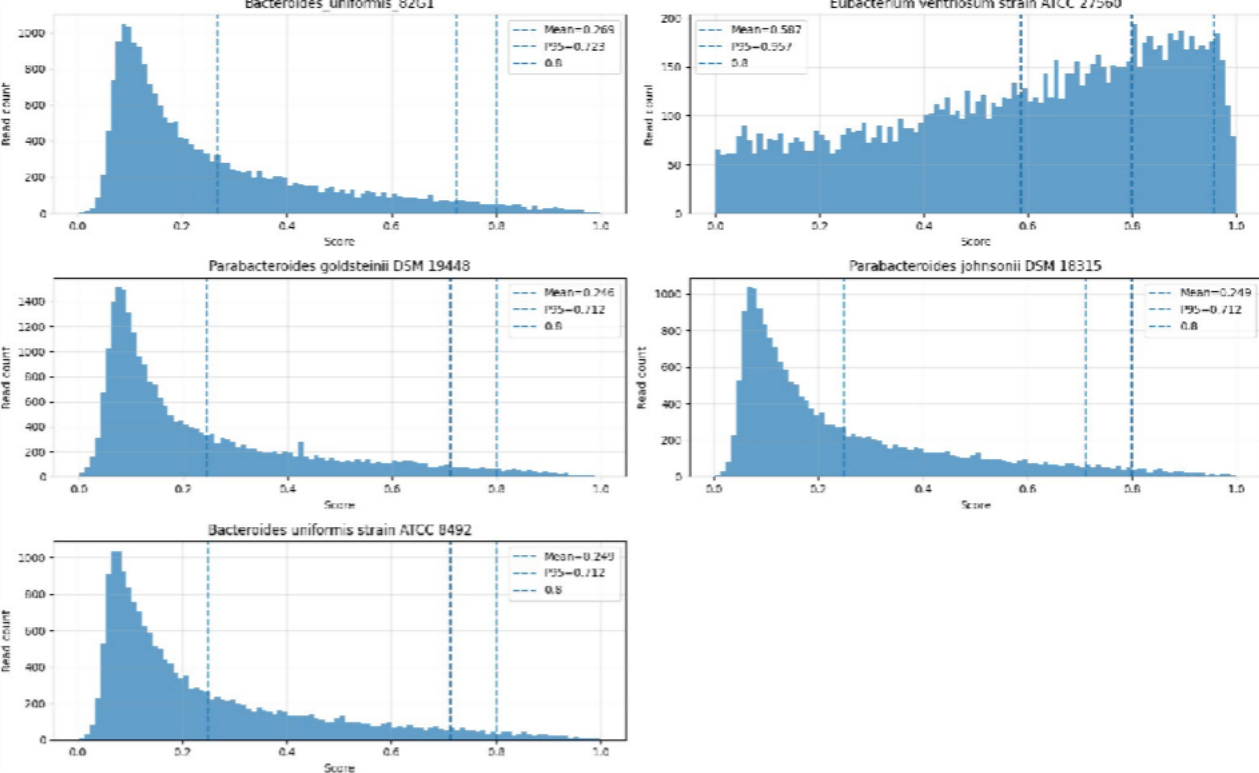

**Figure S6. DeePaC (250 bp) read-level predictions for nonpathogenic genomes.** Four commensal strains show PathFrac values below 0.5 (dashed line), yielding correct nonpathogenic calls. *Eubacterium ventriosum* ATCC 27560 is an outlier, with a score peak near 0.58 and PathFrac = 0.6385 exceeding the 0.5 threshold, driving a false-positive pathogenic prediction. Embedded bar charts show PathFrac (red) versus NonPathFrac (gray).

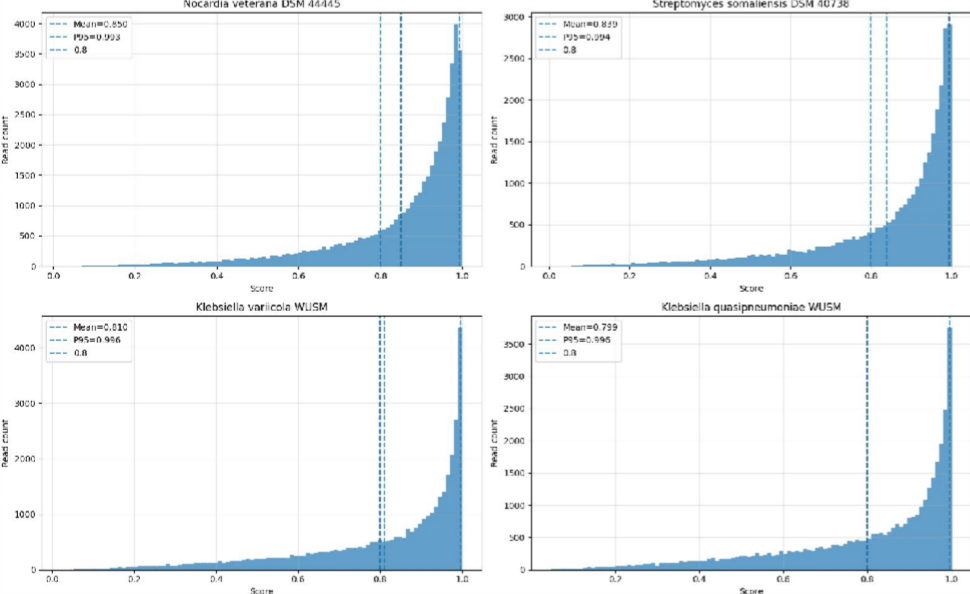

**Figure S7. DeePaC (150 bp) read-level predictions for pathogenic genomes.** For all four pathogenic strains, score distributions are strongly right-skewed with PathFrac values of 0.888–0.944, well above the 0.5 genome-level threshold (dashed line). All strains are correctly classified as pathogenic. Embedded bar charts show PathFrac (red) versus NonPathFrac (gray).

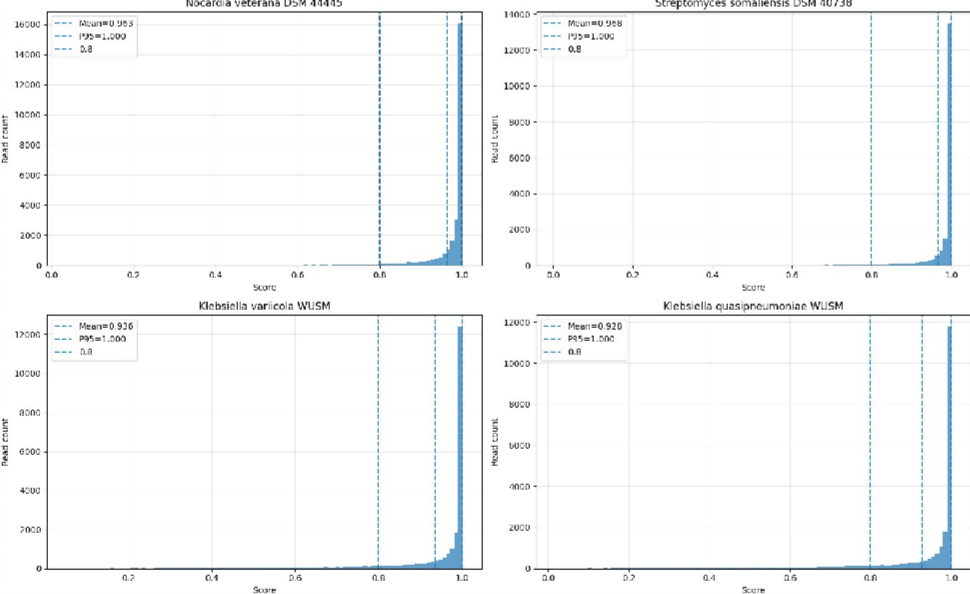

**Figure S8. DeePaC (250 bp) read-level predictions for pathogenic genomes.** All four pathogenic strains show score distributions with prominent peaks at high pathogenicity values (0.58–1.00). PathFrac values range from 0.970 to 0.991, well above the 0.5 genome-level threshold (dashed line), yielding 100% sensitivity. Embedded bar charts show PathFrac (red) versus NonPathFrac (gray).
